# Cross-subject Mapping of Neural Activity with Restricted Boltzmann Machines

**DOI:** 10.1101/2023.04.14.536854

**Authors:** Haoming Yang, Marko Angjelichinoski, Suya Wu, Joy Putney, Simon Sponberg, Vahid Tarokh

**Affiliations:** Department of Electrical and Computer Engineering, Duke University, Durham, NC, USA; School of Physics and Biological Sciences, Georgia Institute of Technology, Atlanta, GA, USA

**Author notes:** Equal Contribution (Haoming Yang, Marko Angjelichinoski). Marko Angjelichinoski and Suya Wu contributed to this work when they were at Duke University. Joy Putney contributed to this work when she was at Georgia Institute of Technology.

## Abstract

Subject-to-subject variability is a common challenge in generalizing neural data models across subjects. While many methods exist that map one subject to another, it remains challenging to combine many subjects in a computationally efficient manner, especially with features that are highly non-linear such as when considering populations of spiking neurons or motor units. Our objective is to transfer data from one or more target subjects to the data space of one or more source subject(s) such that the neural decoder of the source subject can directly decode the target data when the source(s) is not available during test time. We propose to use the Gaussian-Bernoulli Restricted Boltzmann Machine (RBM); once trained over the entire set of subjects, the RBM allows the mapping of target features on source feature spaces using Gibbs sampling. We also consider a novel computationally efficient training technique for RBMs based on the minimization of the Fisher divergence, which allows the gradients of the RBM to be computed in closed form, in contrast to the more traditional contrastive divergence. We apply our methods to decode turning behaviors from a comprehensive spike-resolved motor program – neuromuscular recordings of spike trains from the ten muscles that control wing motion in an agile flying Manduca sexta. The dataset consists of the comprehensive motor program recorded from nine subjects driven by six discrete visual stimuli. The evaluations show that the target features can be decoded using the source classifier with an accuracy of up to 95% when mapped using an RBM trained by Fisher divergence.

**Significant Statement:** In this study, we address the variability of neural data across subjects, which is a significant obstacle in developing models that can generalize across subjects. Our objective is to create a task-specific representation of the target subject signal in the feature space of the source subject. Our proposed RBM architectures achieve highly flexible and accurate cross-subject mapping with few assumptions. Our Fisher RBM improved the previous state of the art method by 300%. Our methods show promise in generalizing features of complex neural datasets across individuals, tuning neural interfaces to subject-specific features, and leveraging data across multiple subjects when experiments are limited in time or completeness.

## 1 Introduction

Combining data across individuals or subjects is a ubiquitous need in analyzing neuroscience experiments. Individual variation can produce subject specific noise and different subjects may utilize different strategies to represent the same sensory input or execute the same motor task. Yet, implementing reliable *cross-subject* algorithms in neuroscience is a notoriously challenging problem. Another important factor contributing to its difficulty arises from the *non-stationary* nature of the neural activity signals, whose statistical properties vary dramatically even under slight changes in the recording conditions (Dabagia et al., 2023; Jayaram et al., 2016; Rao, 2013). As a result, the algorithms trained and optimized on data collected from a given subject, have long failed to perform reliably when directly applied to other subjects (Torres-Oviedo and Ting, 2010). Even modern neural decoders trained on one subject will perform close to a random choice classifier if applied directly to a different subject, thus failing to identify the correct neurological state or stimulus condition even when the subjects perform the same tasks simultaneously (Angjelichinoski, Choi, Banerjee, Pesaran and Tarokh, 2020).

Problems of this type, i.e., problems where the training and test data originate from different distributions, are common in machine learning and are typically tackled within the sub-field of *transfer learning*. Transfer learning methods could have a significant impact on neuroscience and behavior: for example, a reliable cross-neuron population mapping in the latent domain of neural activities can stabilize and improve brain-computer interface (Degenhart et al., 2020; Karpowicz et al., 2022; Pandarinath et al., 2018). In the context of the cross-subject problem, various approaches have already been considered (Jayaram et al., 2016). One direction to map between subjects is domain adaptation methods that map the target data onto the source feature spaces (Angjelichinoski, Choi, Banerjee, Pesaran and Tarokh, 2020; Harvey et al., 2024). Generative modeling is another promising approach to cross-subject mapping: one can apply a directed graph such as the conditional variational autoencoder (cVAE) to generate the target data onto the feature space of the source data (Angjelichinoski, Pesaran and Tarokh, 2020), where the learning model for the downstream task is trained These approaches, however, require a separate directed graphical model to be trained each time a new downstream task and/or new source subject is considered.

A more general approach to cross-subject mapping should be independent of downstream tasks, which are generally tackled through distribution alignment (Dyer et al., 2017; Lee et al., 2019). These works aim to find a mapping that aligns the distribution of target features to the distribution of source features. These methods generally require strong assumptions, which ensures their provably robust, accurate result. For example, Procrustes alignment-based methods require both source and target to exist during test time (Degenhart et al., 2020; Haxby et al., 2011). Another example, the Hierarchical Wasserstein Alignment (HiWA) assumes the alignment can be solved through a linear transformation (Lee et al., 2019), and requires cluster structure in the dataset with cluster labels to solve the optimal transport optimization. These alignment-based methods perform well in many decoding problems where aligning the data of the same subject across time and distributional shifts is needed (Dyer et al., 2017; Karpowicz et al., 2022; Lee et al., 2019). However, in a more complex multi-subject setting where relationships across subjects could be highly non-linear, the strong assumptions of alignment methods restrict their flexibility. These alignment methods could also be inefficient with high-dimensional datasets, as the matrix operations become more computationally intensive as dimensionality grows.

While alignment methods are widely applied in neuroscience research, their restrictions call for reliable non-linear cross-subject mapping methods that are more flexible against complex real world data. One instrumental approach is the Deep Learning method that solves the cross-subject mapping problem through non-linear neural networks. The developed methods applicable to cross-subject mapping usually lie in the field of Transfer Learning (Soleimani and Nazerfard, 2021); however, these methods, especially those based on directed graphs with encoding and decoding processes remain inefficient in the multi-subject scenario (Angjelichinoski, Choi, Banerjee, Pesaran and Tarokh, 2020; Angjelichinoski, Pesaran and Tarokh, 2020). For example, to transfer one subject to 3 different subjects, one would need to train 3 different transfer learning models to achieve a single-direction cross-subject mapping, and the number of models scales exponentially as the number of subjects grows.

In this paper, we propose an efficient, non-linear algorithm that uses undirected graphs to generate samples for cross-subject mapping. Specifically, we leverage the non-linear generative algorithm known as the Restricted Boltzmann Machine (RBM) (Carreira-Perpinan and Hinton, 2005; Hinton, 2002), which allows us to bring the flexible and assumption-less deep learning perspectives and the versatilities of the alignment-based methods together to construct an efficient cross-subject mapping method independent of its downstream decoding tasks RBM is a popular generative model that has had notable success in representation learning with applications in a wide variety of tasks in neuroscience including neuroimaging, classification of temporal events, and EEG analysis (Chai et al., 2017; Hajinoroozi et al., 2016; Kim et al., 2020; Li et al., 2015; Plis et al., 2014). RBMs have also found success in transfer learning, but mainly in computer vision applications (Farahani et al., 2020; Wei and Pal, 2011; Wu and Ji, 2016). The applications of RBM in neuroscience and transfer learning motivate us to adapt RBM for cross-subject mapping, which can serve as a general method with few assumptions on the dataset.

In the neuroscience context, we tackle the general problem of finding a common latent representation of high dimensional spiking data from many neurons across many individual subjects and mapping new subjects to this latent domain. Conceptually what we wish to do is map population recordings from different subjects onto a common latent space and including subjects on which the model has not be previously trained, and then decode from this space. While many other methods have aligned one target to one subject, few have considered the more general problem of combining representations across many subjects even though this is common to many experiments (Hajinoroozi et al., 2016; Kim et al., 2020). Only a few of the current methods consider highly non-linear representations such as spikes but only apply them in a naive environment where only two subjects exist and source and targets both exist during test time (Degenhart et al., 2020; Lee et al., 2019). Here we utilize RBMs to learn a common decoder either from one source subject applied to many test subjects or from a combination of many source subjects applied to a single target. Therefore, during testing, the neural decoder trained on source data can be directly applied to decode the test data of the target subjects because the cross-subject mapping process is independent of downstream tasks. This particular feature of the undirected graphs allows efficient transfer between all subjects with one model, in comparison of the multi-model method induced by traditional subject mapping One limitation of RBMs is that they often are computationally expensive because contrastive divergence minimization does not have a closed form and requires Gibbs sampling. Here, we also extend the conventional contrastive divergence training of RBM (which is equivalent to maximum likelihood, we refer more details to (Hinton, 2002)) by proposing an alternative training based on Fisher divergence minimization (Hyvärinen, 2005). The Fisher divergence minimization allows the gradient of the RBM to be computed in closed form, fostering an even more efficient implementation that does not require iterative Gibbs sampling during training.

We evaluate the performance of our method for cross-subject decoding of discrete visual stimulus conditions using the spiking activity of the motor program, specifically the set of spiking motor units, in nine hawk moths (Putney et al., 2021). Each moth is exposed to the same set of six visual stimuli and the neuromuscular activity is collected in the form of spike trains extracted from fine wire electromyography (EMGs) of the ten primary flight muscles that control the wings, resulting in a comprehensive, spike-resolved motor program (Putney et al., 2019). Unlike vertebrate EMGs, these flight muscles act as effectively single motor units and result in identifiable spike trains comparable to population recordings of individual units elsewhere in the brain or peripheral nervous system.

In the context of these specific data, the challenge is to enable the decoding of a new target subject moth’s motor program into the motor outcome (turn direction) based on a set of latent space shared in common with the other subjects (i.e. individual moths). Within a single subject, we can already nearly perfectly decode behavior as expected from a comprehensive motor program recording. However, the latent structures within population recordings across the different moths are different, so naive decoding does not generalize. Using our RBM with Fisher divergence we show that we can learn a generalized latent domain from the population of spiking units. We compare the decoding accuracy of this method to other common transfer learning methods most notably HiWA and RBMs with more traditional contrastive divergence. Our results demonstrate the promising potential of the proposed framework, with respectively up to 90% and 95% accuracy in decoding the behavioral state (i.e., the visual stimulus), when using the RBM trained with classical and the new Fisher divergence-based methods; these results thoroughly improve from previous state-of-the-art distribution alignment-based methods like HiWA.

## 2 Material and Methods

We divide this section into three parts. In Section 2.1 we present the statistical formulation of the problem of cross-subject mapping as a problem of learning joint distribution between target and source feature vectors. Next, in Section 2.2, we discuss RBMs and present both the contrastive divergence and Fisher divergence training methods. Section 2.3 presents a simple cross-subject mapping approach that uses Gibbs sampling to draw samples from the joint distribution of the target and source features. Our code to replicate our study is available in the extended data.

### 2.1 Problem Statement

The objective of cross-subject learning is to map target subject features to the feature space of source subjects so that the motor intentions of the target subject can be decoded by the neural decoder that is learned on source subjects. In other words, we aim to obtain the appropriate source space representation of the tasks that the target subjects perform. Technically, the problem boils down to finding a function that maps the feature vectors. This mapping function can be assumed to be purely deterministic. However, in this paper, we adopt a probabilistic approach, which generates task-specific features in source feature space. We outline details below.

Let *ℳ*_D_ and *ℳ*_S_ denote the index sets of the target and source subjects, respectively. Source subject(s) is defined as the subject(s) where the neural decoder is available during test time, and target subject(s) is the subject that we wish to decode using the source decoder Further, we let *ℳ* denote the joint set of all subjects, namely *ℳ* = *ℳ*_D_ ∪*ℳ*_S_. For simplicity, we assume that the subsets *ℳ*_D_ and *ℳ*_S_ are disjoint, namely 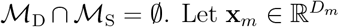 denote the *D*_*m*_-dimensional vector representing the neural activity of subject *m* ∈ *ℳ*; we refer to **x**_*m*_ as the *feature vector* of subject *m*. Furthermore, we use 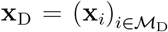 and 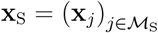 to respectively denote the joint feature vectors of the target and source subjects. Note that **x**_D_ and **x**_S_ are vectors with dimensions 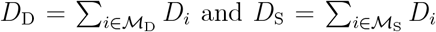, respectively. Finally, we use **x** to denote the joint vector of features of *all* subjects in the set *ℳ*; that is **x** = (**x**_D_, **x**_S_) = (**x**_*m*_)_*m*∈*ℳ*_, and its dimension *D* = _*m*∈*ℳ*_ *D*_*m*_.

To learn the cross-subject mapping, we consider a *conditional probability distribution* to generate feature representations in the feature space of source subjects given the feature vector of target subjects. One option is to directly parameterize the probability density function *p*(**x**_S_|**x**_D_) of this conditional distribution. Another approach is to first learn the distribution *p*(**x**_D_) and the joint distribution *p*(**x**_S_, **x**_D_) of all feature vectors across the entire population of subjects in *ℳ*, and then to obtain the conditional distribution *p*(**x**_S_|**x**_D_) by Bayes’ theorem.

For the purpose of cross-subject mapping, it is not mandatory to learn an explicit probability density function (pdf) of the conditional distribution. Recall that the objective is to obtain the feature representations of the target subject in the feature space of the source subject. Therefore, to this end, we aim to learn a generative framework such that we can easily sample from the conditional distributions *p*(**x**|**h**; *θ*) and *p*(**h**|**x**; *θ*). Generative algorithm for cross subject mapping in previous works, generally applies directed graphs such as cVAE (Angjelichinoski, Pesaran and Tarokh, 2020; Sohn et al., 2015), which map the target feature onto the feature space of the source subjects by the decoder of the autoencoder. However, directed graphs are not flexible to adopt new source subjects, and the architecture of cVAE is built on deep neural networks which are not easy to fine-tune given the limited size of data. In this paper, we consider the class of undirected graphical models and easily adapt to the generative framework. Given its relatively simple architecture and straightforward sampling scheme, an RBM is the first generative model we will tackle. In our application, the source data is unknown during test time, and our following construction of the RBM allows the mapping of multiple subjects from targets to sources with only one trained model.

This RBM is constructed such that a joint distribution of all features is learned. Specifically, we consider a generative model *p*(**x**) of the concatenated feature vector **x** = (**x**_D_, **x**_S_). We first learn the joint distribution *p*(**x**) = *p*(**x**_D_, **x**_S_). Once the joint distribution is learned by the RBM, we use Gibbs sampling to sample from *p*(**x**) with hidden variables **h**. The Gibbs sampler proceeds as follows: we initialize the visible variables by 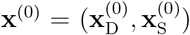. Here, 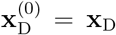 is given, and 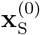 is a dummy and noise-like vector. Next, we sample hidden variables **ĥ** from *p*(**h**|**x**; *θ*), and then sample visible variables **x**^(1)^ from *p*(**x**|**h**; *θ*). After sufficient sampling iterations, we expect to obtain 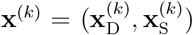 where the 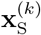 is the feature representation of **x**_D_ in the feature space of source subjects. The hidden layer **h** is designed to bridge target and source feature vectors. Note that this architecture setup allows us to only train one RBM that covers the need for any multi-subject mapping during test time. This flexibility follows from learning the joint distribution during training, allowing us to efficiently apply Bayes rule during sampling. During the training of the RBM, there is no distinction between **x**_D_ and **x**_S_ as all subjects are available. During test time however, only *ℳ*_S_’s neural decoder and only *ℳ*_D_’s data is available. With reliable conditional sampling from the learned conditional distribution, we can now map new data from target to source without any source data available during testing, and apply the sampled data to the source’s neural decoder. We illustrate this generative framework in Fig. 1.

**Fig 1:**
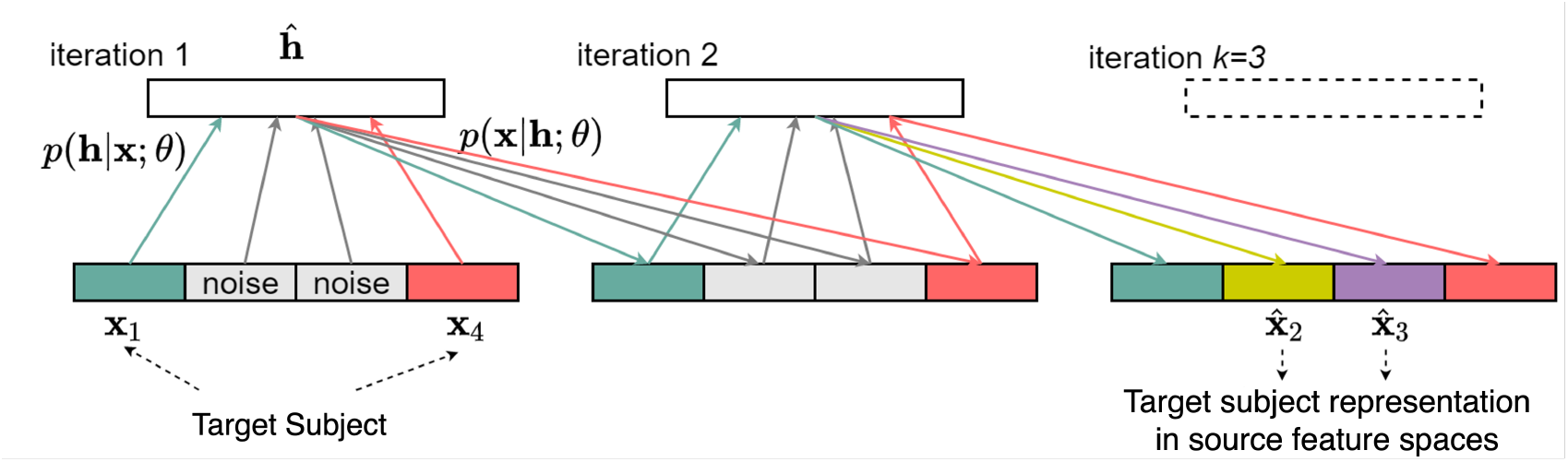
A generative framework for the cross-subject mapping during testing. We assume that both the source and target data are available during training. Neural decoders, which in our case are linear classifiers (see details in section 3), are trained for the source subjects. Only the target subjects are available during testing, and we replace the source representation with randomly sampled standard Gaussian noise. The trained RBM, after *k* iterations, will sample from the conditional distribution *p*(**x**_S_ **x**_D_), which can be seen as the target representation in source space. In this figure *k* = 3.

### 2.2 Learning Restricted Boltzmann Machines

We will first discuss the Gauss-Bernoulli RBMs and outline the principles of their training. We then review the standard training technique that aims to minimize the contrastive divergence, which is equivalent to minimizing the Kullback-Leibler (KL) divergence between the data-generating distribution and the model (Carreira-Perpinan and Hinton, 2005; Hinton, 2002). We also consider an alternative training technique that minimizes the Fisher divergence (Hyvärinen, 2005). Unlike the classical method, minimizing the Fisher divergence approach allows the gradients of the loss function to be computed in closed form. This improves the computational efficiency and reliability of the training.

#### 2.2.1 Notation

We adopt the following notation conventions. Recall that in the context of the cross-subject mapping problem outlined in Section 2.1, the vector **x** comprises the feature vectors of the entire population of subjects, i.e., **x** = (**x**_1_, …, **x**_*M*_). Let ∇_**x**_ and Δ_**x**_ denote the gradient and Laplacian operator with respective to (w.r.t.) the vector **x**. Let dg(**x**) denote a diagonal matrix whose main diagonal is **x**. For a square matrix **A**, let dg(**A**) denote a diagonal matrix formed by setting all the elements to **A** not on the main diagonal to zeroes. We use ∥**x**∥ (respectively ∥**A**∥) to denote the *L*_2_-norm of the **x** (respectively the Frobenius norm of **A**). We further use *p*_∗_(**x**) to denote the true data-generating distribution of **x**. In practice, *p*_∗_(**x**) is usually unknown; therefore, given a set of observations, a standard problem is to estimate the model density *p*(**x**) from some model class that best explains the data under an appropriate evaluation metric. In this paper, we focus on a parametric density model class *p*(**x**; ***θ***), ***θ*** ∈ Θ, which is parameterized as an RBM (see Section 2.2.4).

#### 2.2.2 Kullback-Leibler Divergence and the Logarithmic Loss

A common practice to measure the deviation of a postulated probability distribution *p*(**x**) from the data-generating distribution *p*_∗_(**x**) is to use the KL divergence defined by

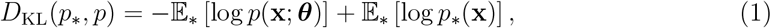

where the expectation is taken w.r.t. *p*_∗_(**x**) (as denoted by the subscript). *D*_KL_(*p*_∗_, *p*) ≥ 0 with equality if and only if *p* = *p*_∗_ almost surely. The minimization of *D*_KL_(*p*_∗_, *p*) is equivalent to the minimization 𝔼_∗_ [*ℓ*(**x**; ***θ***)] where *ℓ*(**x**; ***θ***) = − log *p*(**x**; ***θ***) is called the logarithmic loss. Let ***θ***_∗_ denote the data-generating parameter that minimizes the KL divergence in Eq.(1), namely *p*(**x**; ***θ***_∗_) is closest to *p*_∗_(**x**) among all distributions over Θ under the KL divergence. It can be shown from the law of large numbers and standard regularity conditions (White, 1982) that the maximum likelihood estimate (MLE) 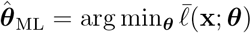 satisfies 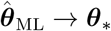 in probability as the number of data points grows. In other words, the MLE is consistent.

#### 2.2.3 Fisher Divergence and Hyvärinen Score

The Fisher divergence of *p*(**x, *θ***) from the data-generating pdf *p*_∗_(**x**) is defined by

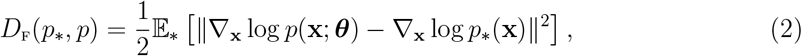

where the expectation is again taken w.r.t. the data-generating pdf *p*_∗_(**x**). We note that *D*_F_(*p*_∗_, *p*) ≥ 0 with equality if and only if *p*_∗_ = *p* almost surely. Under mild regularity conditions, the Fisher divergence Eq.(2) can be written as (Hyvarinen, 2005)

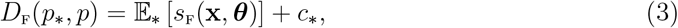

where *c*_∗_ is a term that does not depend on ***θ*** and *s*_F_(**x, *θ***) is the Hyvärinen Score, defined as

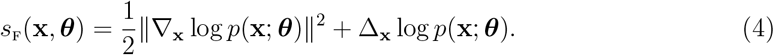

The result Eq.(3) enables to minimize the Fisher divergence over the space of parameters Θ by minimizing the empirical analog of the Hyvärinen Score 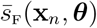. Let ***θ***_∗_ denote the parameter value that minimizes the Fisher divergence between *p*(**x**; ***θ***_∗_) to *p*^∗^(**x**) between all model class candidates. By standard asymptotic analysis, it can be shown that the estimate 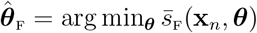 satisfies 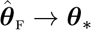 in probability as the number of data points grows (we refer details to (Hyvärinen, 2005) and references therein). This estimation procedure is known as score matching. It has been proved that score matching using the Langevin Monte Carlo method is equivalent to contrastive divergence in the limit of infinitesimal step size (Hyvärinen, 2007). Although this result implies that this variant of convergence divergence can retain the consistency on score matching, we note that this equivalence holds only for a particular MCMC method. The actual performance of these two methods are different.

#### 2.2.4 Gauss-Bernoulli Restricted Boltzmann Machines

An RBM is a bipartite undirected graphical model where only the links between visible units and hidden units are allowed. We focus on Gauss-Bernoulli RBM, which consists of continuous inputs **x** ∈ ℝ^*D*^ and binary hidden units **h** ∈ {0, 1}^*M*^ with the pdf

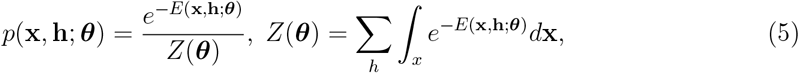

where the energy function *E*(**x, h**; ***θ***) is given by

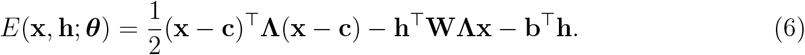

Here, **W** ∈ ℝ^*H*×*D*^ is the matrix of weights connecting the units from the hidden and input layer, **b** ∈ ℝ^*H*^ and **c** ∈ ℝ^*D*^ are the vectors of hidden and input layer biases, and **Λ** = dg(***λ***) denotes the diagonal precision matrix of the inputs. All these parameters are freely learnable and they are denoted by ***θ*** in Eq.(5) and Eq.(6). We illustrate a Gauss-Bernoulli RBM model in Fig. 2. It is easy to see that the conditional densities are given by

**Fig 2:**
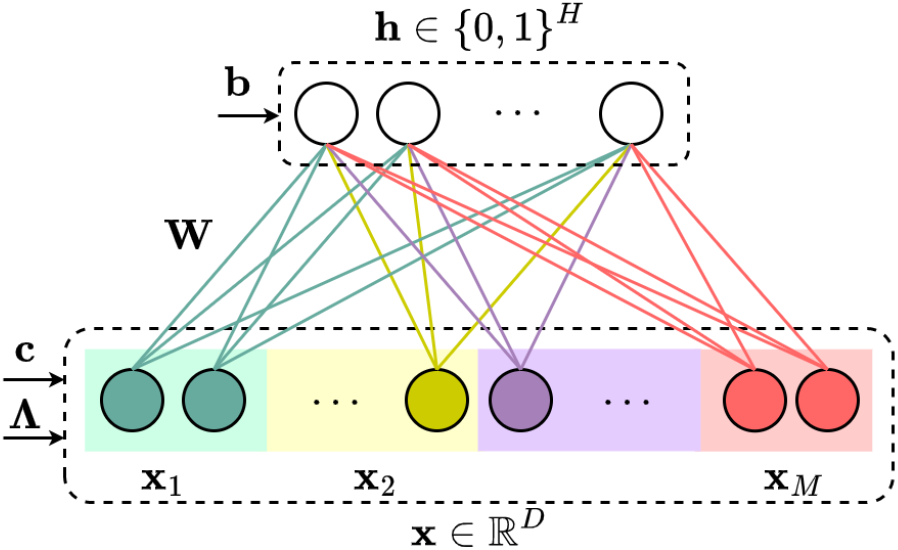
A bipartite undirected graphical model: Gauss-Bernoulli restricted Boltzmann machine with continuous input and binary hidden units.

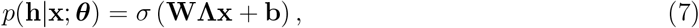

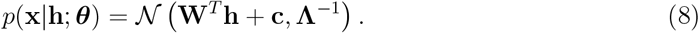

The marginal density *p*(**x**; ***θ***) of the visible inputs can be also written in the energy-based form

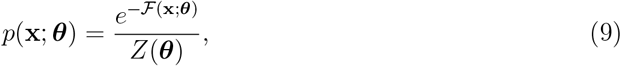

where *Z*(***θ***) is called the normalizing constant, and ℱ(**x**; ***θ***) is the free energy:

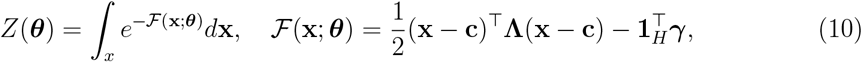

with ***γ*** = log(**1**_*H*_ + exp (**WΛx** + **b**)) denoting the element-wise Softplus function. Unlike Eq.(6), the energy function Eq.(10) associated with the marginal *p*(**x**; ***θ***) is no longer linear in the free parameters ***θ***.

A frequently encountered Gauss-Bernoulli RBM in the literature is the one associated with the conditional density *p*(**x**|**h**; ***θ***) = 𝒩 (**W**^*T*^ **h** + **c, I**_*D*_) and assumes unit variances for the input units. This is a special case of our model Eq.(6) in which we treat the variances of the inputs as learnable parameters, and all results and discussions in this paper can be applied in a straightforward manner to the special case by replacing **Λ** with the identity matrix.

#### 2.2.5 Learning RBM via Contrastive Divergence

The negative log-likelihood of the parameters of the RBM, i.e., the logarithmic loss can be written as

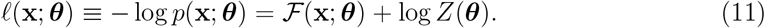

The gradient obtains a particularly interesting form:

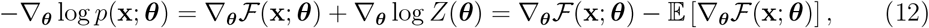

where the expectation in the second term is taken w.r.t. the marginal density of the visible units given in Eq.(9). Therefore, it is difficult to determine the gradient analytically. In order to make the computation tractable, this expectation is estimated using samples from *p*(**x**; ***θ***) which can be obtained by running a Markov chain with Gibbs sampling as the intermediate sampling operator. To speed up the sampling process, Hinton (2002) showed that it is not necessary to wait for the Markov chain to converge; instead, if the chain is initialized using training examples, reasonable learning performance might be obtained only after *k* Gibbs steps. In practice, *k* = 1 is commonly used during training (Hinton, 2002). However, this corresponds to the approximate minimization of the contrastive divergence (CD), which produces biased estimates of the model parameters (Carreira-Perpinan and Hinton, 2005). While obtaining an expected value may require a large *k* for a contrastive divergence to converge (van der Plas et al., 2023), in our application, we need to sample from the conditional distribution, therefore we chose *k* = 1 during test time where we sample from the distribution of source conditioned on target

We see that an important implication of approximating MLE-based learning through contrastive divergence minimization is the lack of consistency guarantees. Specifically, minimizing the contrastive divergence is not guaranteed to converge to the data-generating parameter ***θ***_∗_ that minimizes the KL divergence from *p*(**x**; ***θ***) to the data-generating pdf *p*_∗_(**x**). The impediment can be traced back to the computation of the gradient of the logarithmic loss and the analytical intractability of the second term in Eq.(12) which appears due to the intractability of the partition function as a normalizing constant in Eq.(5).

#### 2.2.6 Learning RBM via Fisher Divergence

To overcome the issues associated with the lack of consistency guarantees, instead of aiming to minimize the KL divergence through contrastive divergence approximation, we propose to minimize the Fisher divergence from the marginal density of the visible units *p*(**x**; ***θ***) to the data-generating distribution *p*_∗_(**x**). To evaluate the Hyvärinen Score Eq.(4) based on Eq.(9), we derived the following result.

**Proposition 1** *The Hyvärinen Score for the Gauss-Bernoulli RBM Eq*.(5) *with energy function Eq*.(6) *is given by*

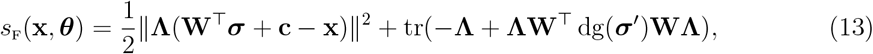

*with* ***σ*** = *σ*(**WΛx** + **b**) *and* ***σ***^′^ = *σ*^′^(**WΛx** + **b**), *where σ and σ*^′^ *respectively denote the element-wise Sigmoid operator and the corresponding first derivative*.

**Proof 1** *Taking the derivative of the log-density* log *p*(**x**; ***θ***) *w.r.t*. **x**, *we obtain*

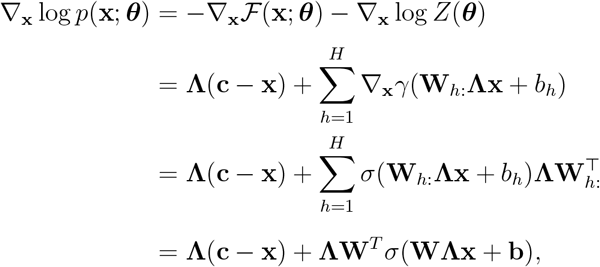

*which gives the first term in Eq.(13)*. *To obtain the second term, we first compute the Hessian matrix of the log-density* log *p*(**x**; ***θ***); *we obtain:*

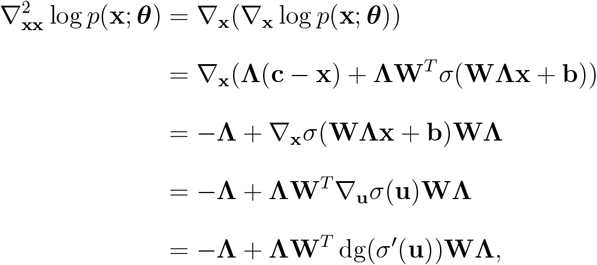

*where* **u** = **Wx**+**b**. *Plugging the Hessian into the Laplacian* 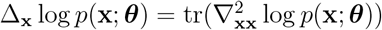 *gives the second term in Eq*.(13), *which completes the proof*.

We observe that unlike the logarithmic loss in Eq.(11), the Hyvärinen Score can be evaluated explicitly in terms of the parameters **c, b** and **W** of the Gauss-Bernoulli RBM. Moreover, the calculation does not involve the partition function *Z*(***θ***). This simplifies the computation of the gradient w.r.t. the parameters of the RBM, which now can be computed by straightforward application of matrix calculus yielding the following closed-form expressions:

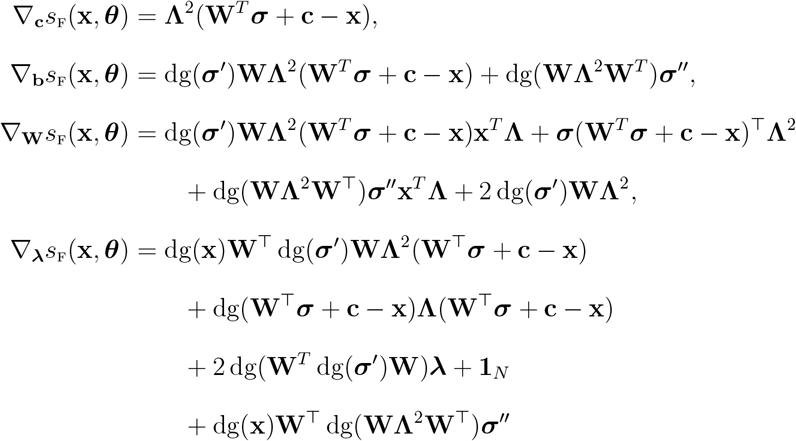

with ***σ*** = *σ*(**WΛx** +**b**), ***σ***^′^ = *σ*^′^(**WΛx** +**b**) (as in Proposition 1), whereas ***σ***^′′^ = *σ′′* (**WΛx** +**b**); also, recall that **Λ** = dg(***λ***).^1^

It is evident that, as opposed to the gradient of the logarithmic loss Eq.(12), the gradient of the Hyvärinen Score can be computed explicitly w.r.t. the parameters of the Gauss-Bernoulli RBM, producing closed-form expressions that can be used directly used for training the parameters of the RBM. Indeed, let 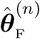 denote the parameter estimate at each step *n*. The new parameter estimate can be obtained through the following update rule (repeated until convergence):

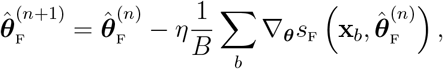

where *η* is the learning rate, *B* is the size of the minibatch of randomly chosen data points **x**_*b*_, and *b* = 1, …, *B* at step *n*.

### 2.3 Cross-subject Mapping Algorithm

Recall from Section 2.1 that in cross-subject mapping, the goal is to obtain source feature representation(s) **x**_S_ from the target feature vector(s) **x**_D_. We will elaborate on how to use an RBM and the Gibbs sampler to sample such representations. We first parameterize the generative model *p*(**x, h**; ***θ***) of all feature vectors **x** = (**x**_S_, **x**_D_) and hidden variables **h** using the Gauss-Bernoulli RBM described in Section 2.2.4, see also Fig. 2. After learning the parameters of the Gauss-Bernoulli model we infer 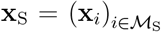 from 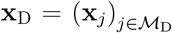 as follows. First, we initialize the vector 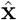 by the features of the target subjects **x**_*m*_, *m* ∈ *ℳ*_D_ and random noise (e.g., with standard normal variables). Then:

1. generate 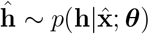 via Eq.(7);
2. using **ĥ** generate 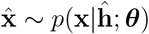 via Eq.(8).

We obtain the final estimate after repeating the above two steps *k* ≥ 1 times; Fig. 1 illustrates an example with *k* = 3. This gives the source feature space representations of the target feature vectors and they can be further processed using algorithms trained on source data. Alternatively, in the final step, we can skip sampling from *p*(**x**|**ĥ**; ***θ***) and we can also infer **x**_S_ as 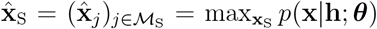. For simplicity and without loss of generality, we have assumed that target and source subjects *ℳ*_D_ and *ℳ*_S_ satisfy *ℳ* = *ℳ*_D_ ∪ *ℳ*_S_ and *ℳ*_D_ ∩ *ℳ*_S_ = ∅. This allows us to skip a tedious step in the algorithm and avoid the marginalization of the joint density *p*(**x, h**; ***θ***) over subjects that are neither targets nor sources.

## 3. Evaluation and Results

Next, we present the results from the evaluations. First, in Section 3.1 we describe the experimental protocol and the acquired data. In Section 3.2 we discuss the evaluation methodology, including evaluation scenarios. In Section 3.3 we present the main findings and observations.

### 3.1 Experiment, Data and Features

#### 3.1.1 Protocol

We study a comprehensive flight motor program for hawk moths. We will describe the experimental protocol and related procedures only briefly here; the interested reader is referred to (Putney et al., 2021) where the data set was first published for more details. The subjects, i.e., the moths are tethered inside a three-sided box formed by computer monitors displaying the visual stimuli. Each stimulus is represented by sinusoidal gratings with a spatial frequency of 20° per cycle on 3D spheres projected on the monitors. The spheres drift at a constant velocity of 100° per second, corresponding to a temporal frequency of 5 cycles/second. Moreover, the spheres also drift in opposite directions about the three axes of rotation which result in 6 different visual stimuli also known as pitch (up, down), roll (left, right), and yaw (left, right) (Putney et al., 2021).

The moth responds to each of the 6 discrete stimuli by producing turning effort as assessed with a 6 degree-of-freedom force/torque transducer. The 10 primary muscles that control the flying motion of the moth are wired and enable spike-resolved EMG signals to be recorded during tethered flight. These key muscles include the flight power muscles (dorsolongitudinal (DLM) and dorsoventral (DVM) muscles), as well as the steering muscles, (third axillary (3AX), basalar (BA), and subalar (SA) muscles) on both the left and the right side of the thorax. The EMG recordings are used to extract the timings of the motor unit spikes in each of the muscles that serve as control commands by means of which the nervous system guides the motion of the moth in response to the different visual stimuli; more details can be found in Putney et al. (2019) and Putney et al. (2021). Taken together, this dataset is unusual in its near-complete recording of all the spikes the animal can use to control its wings and so is an ideal point of convergence to test for the decodability of stimulus conditions.

The objective is to decode the visual stimulus from the comprehensive motor program recordings, i.e., the spike trains. The subject-specific formulation of the problem where the neural decoder (classifier) is both trained and tested on the same subject was analyzed in Putney et al. (2021). Here, we study the performance of the neural decoder in cross-subject settings, where the test data originates from the target subjects whereas the classifier is trained on source subject data.

The dataset is collected from 9 subjects over 20 seconds of recording sessions for each visual stimulus. Each session is segmented into *wing strokes*, i.e., trials (Putney et al., 2021).

The typical duration of a wing stroke is between 50 and 70 milliseconds yielding an average number of trials of ≈ 2500 per moth. It should be noted that some moths in the dataset are missing the recordings from some of their muscles (either one or at most two) due to failures in the recording procedure. Nevertheless, as demonstrated in Putney et al. (2021), the absence of some (one or two) muscles does not have a significant effect on decoding performance; in fact, as shown in Putney et al. (2021), high decoding accuracy (higher than 90%) can be achieved even with half of the available muscles due to the completeness of the motor program.

#### 3.1.2 Feature Extraction

Before we delve into more details with respect to the cross-subject neural decoder, we briefly describe our methodology for constructing feature representations from spike trains proposed in Putney et al. (2021). Since the spike trains are given by variable-length vectors of spike timings, we consider Gaussian kernels, a strategy commonly used in neuroscience, to interpolate the spike trains. The Gaussian kernel is given by:

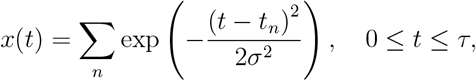

where *t*_*n*_ denotes the timing of the *n*-th spike collected from an arbitrary muscle, trial, and moth, *τ* denotes the wing stroke cut-off threshold (we only consider spikes that satisfy *t*_*n*_ ≤ *τ*), and *σ* is the Gaussian kernel bandwidth. The goal is to obtain a smooth multivariate time-series representation of the spike trains in which the spike timing information is conveyed by centering one kernel at each spike and summing the kernels; these yields feature vectors of fixed dimension *τ* · *ν*_*S*_ where *ν*_*S*_ is the sampling frequency. For consistency, the muscles whose recordings are missing are filled with zero vectors of the same length as above. We then flatten the interpolated time series across muscles to obtain one large feature vector. Finally, we apply PCA and retain only the first *P* largest modes; this is our final representation **x**_*m*_ of the neural activity of subject *m* with dimension *D*_*m*_ = *P*.

Fig. 3 depicts the feature spaces of each moth and all six visual conditions (tasks) across the first two strongest principal components, i.e., modes after performing PCA. The diagrams also show the confidence ellipses corresponding to one standard deviation for each of the conditions. It can be clearly observed that the data demonstrates strong separability properties even in the first two dimensions of the PCA-based feature space; adding more features (i.e., PCA modes) only increases this separability in the higher-dimensional feature space for each moth as we have observed in past work (Putney et al., 2021); for more details, we advise the interested reader to refer to Putney et al. (2021) where the feature extraction procedure was first proposed and its performance thoroughly analyzed.

**Fig 3:**
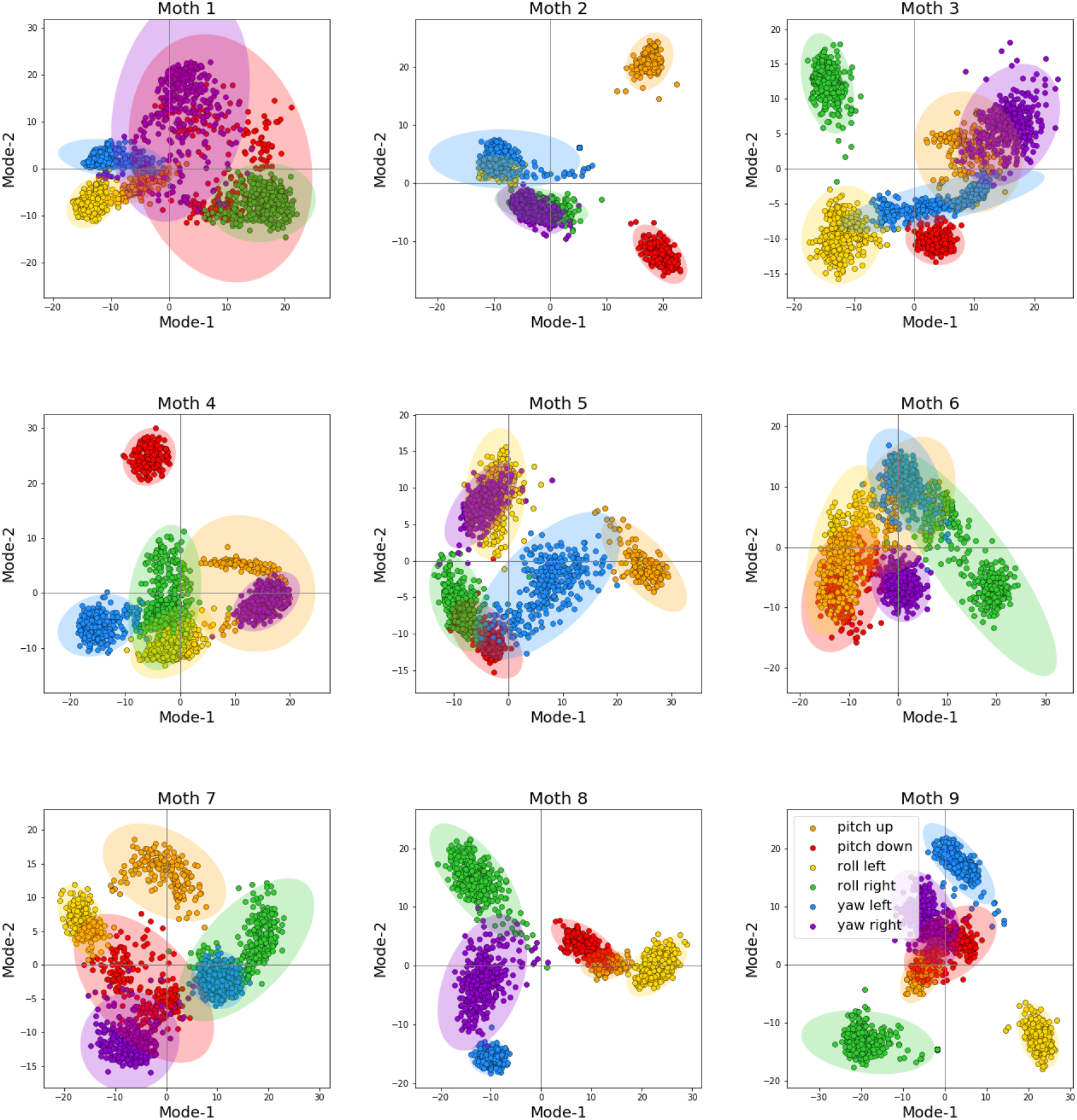
The feature space across the first two principal modes for each moth. For visual clarity, the diagrams also show the confidence ellipse corresponding to one standard deviation for each of the visual conditions.

### 3.2 Scenarios and Methodology

The moth population set is *ℳ* = {1, …, 9}. We evaluate the performance of the cross-subject neural decoder in the following scenarios:

- *Scenario I* (Fig. 4a). The source and target sets are *ℳ*_S_ = {*m*} and *ℳ*_D_ = *ℳ*\{*m*}, respectively. That is, we select a single subject *m* ∈ *ℳ* as the source and map the features of *all* remaining subjects *i*≠ *m* onto the feature space of the source subject *m*. The obtained representation is then decoded using a linear classifier trained on subject *m* data.

**Fig 4:**
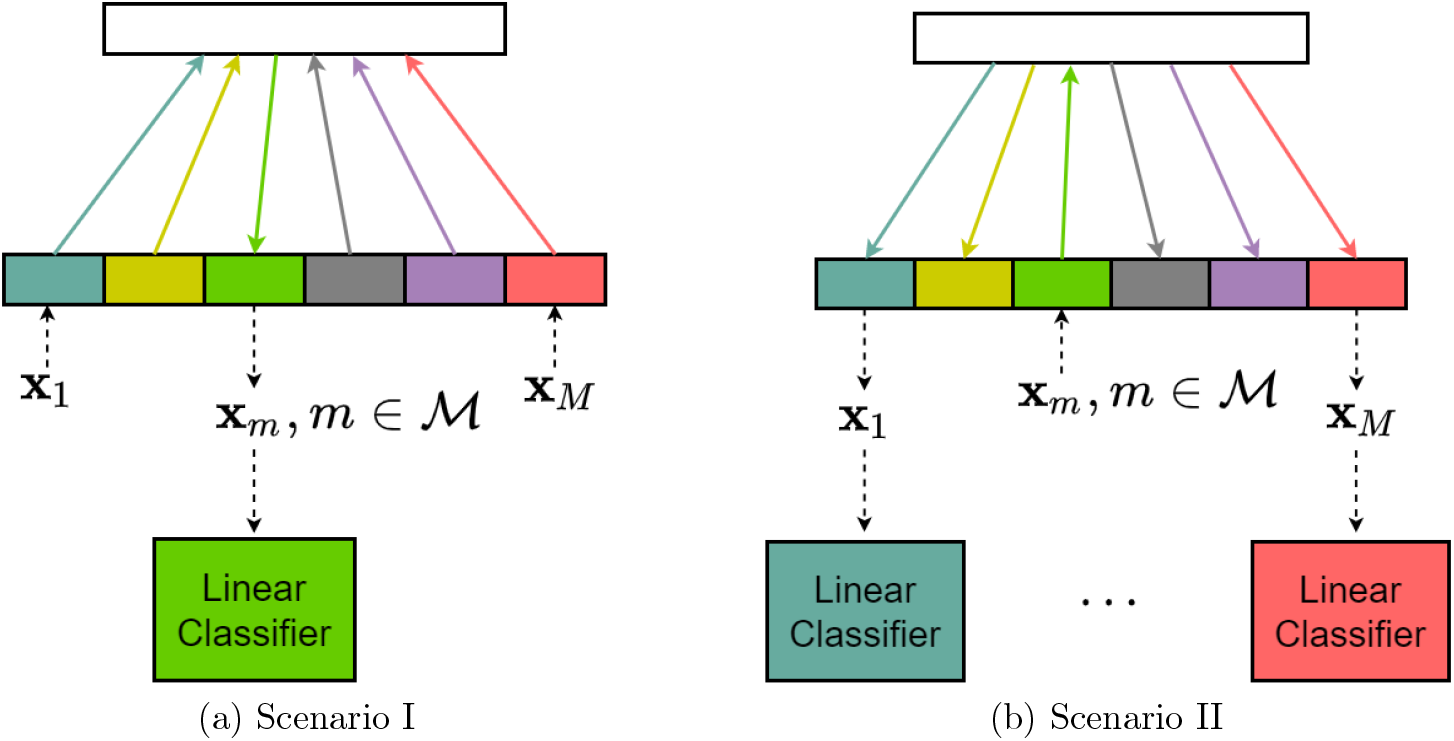
Evaluation scenarios. (a) One subject is the source, all other subjects are targets. (b) One subject is the target, and all other subjects are sources.
- *Scenario II* (Fig. 4b). The target and source index sets are *ℳ*_D_ = {*m*} and *ℳ*_S_ = *ℳ*\{*m*}, respectively. In other words, we select a single subject *m* ∈ *ℳ* as the target and map the corresponding feature vector onto the feature spaces of *all* remaining subjects *j*≠ *m*. Similarly, as in scenario I above, the obtained representations of the target feature vector are subsequently decoded using the subject-specific linear classifiers trained on the source subjects *j*≠ *m* individually.

In both of these scenarios, we evaluate the performance of the source classifier (trained purely on source data) on the transferred target features through an RBM model with both standard contrastive divergence minimization and Fisher divergence minimization; we use RBM-CD and RBM-FD to denote these two RBM models, with CD standing for contrastive divergence (see Section 2.2.5) and FD standing for Fisher divergence (see Section 2.2.6). We compare the performance with three benchmarks:

- subject-specific neural decoding, when the classifier is trained *and* tested on purely source data. Equivalently, the data used to train the classifier and the test data respectively come from the same individual subject. In this case, the performance of the neural decoder may be thought of as an upper bound because it represents the best-case scenario where the classifier has access to the most relevant information about feature patterns of the test data.
- cross-subject neural decoding with *no* transfer, when the target data is directly decoded using the source neural decoder without transferring the target data into source feature space. As we discussed in Section 1, in this case, the performance of the neural decoder is usually close to a random guess, since the neural decoder does not take into account the differences between neural activity patterns between the test (target) and the train (source) data.
- cross-subject neural decoding using HiWA, as described in Section 1, is a supervised state-of-the-art linear alignment technique that exploits the cluster structure to solve a hierarchical optimal transport optimization (Lee et al., 2019). This method is widely applied in neural settings (Dabagia et al., 2023).

The training and testing data for the target and source data sets in both scenarios are formed by splitting the original data sets of each moth into training and testing data subsets randomly according to *ω* ∈ (0, 1), which denotes the ratio between the number of training samples and the total number of trials. To characterize the performance of the cross-subject neural decoder statistically, the random train/test data splitting procedure is repeated 100 times; the entire model including the RBM is then re-trained using the new training data, and the test performance is recorded.

In cross-subject mapping, we are aiming to find a source space feature representation of the target feature vector; the mapping should be consistent with the task that the target features are encoding. In other words, the mapping should be such that the transferred representation of the target feature vector should be in the region of the source feature space that corresponds to the task that the target is performing, as illustrated in the top row in Fig. 5. In essence, when sampling from the joint pdf *p*(**x, h**), we wish to sample from a stimulus-dependent distribution and this should be guided by the task information the target feature vector is encoding. Hence, the trials in the training data set should be organized such that there is an input-output correspondence with respect to the stimulus. As we are unable to establish correspondence between individual training trials, we assign the correspondences at random. The procedure is schematically depicted in the bottom row in Fig. 5. Namely, for each target feature vector from the training set, we randomly choose a source feature vector from the same stimulus class and declare the pair to be an input-output pair.

**Fig 5:**
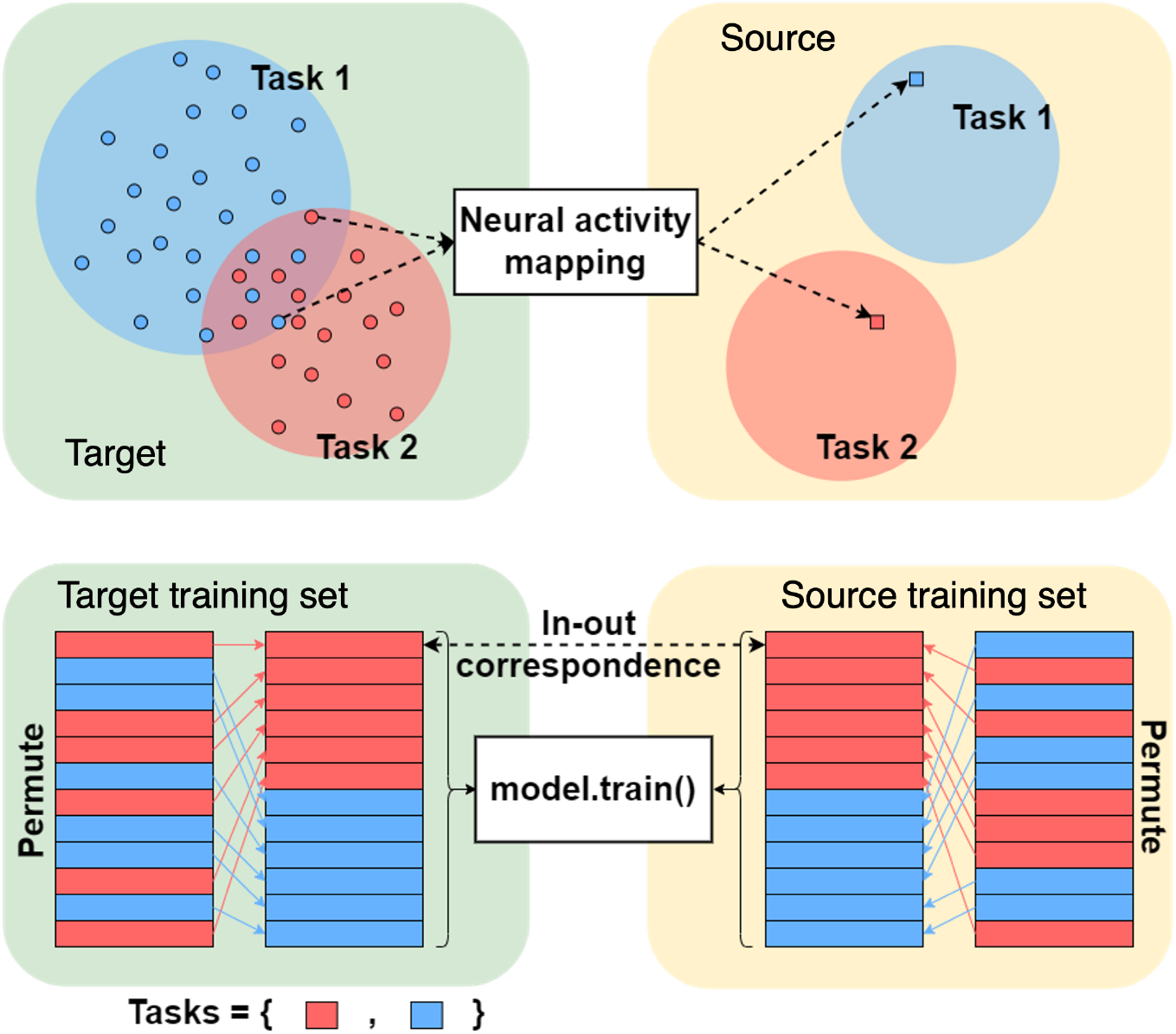
Organization of target and source training data sets.

**Table 1:**
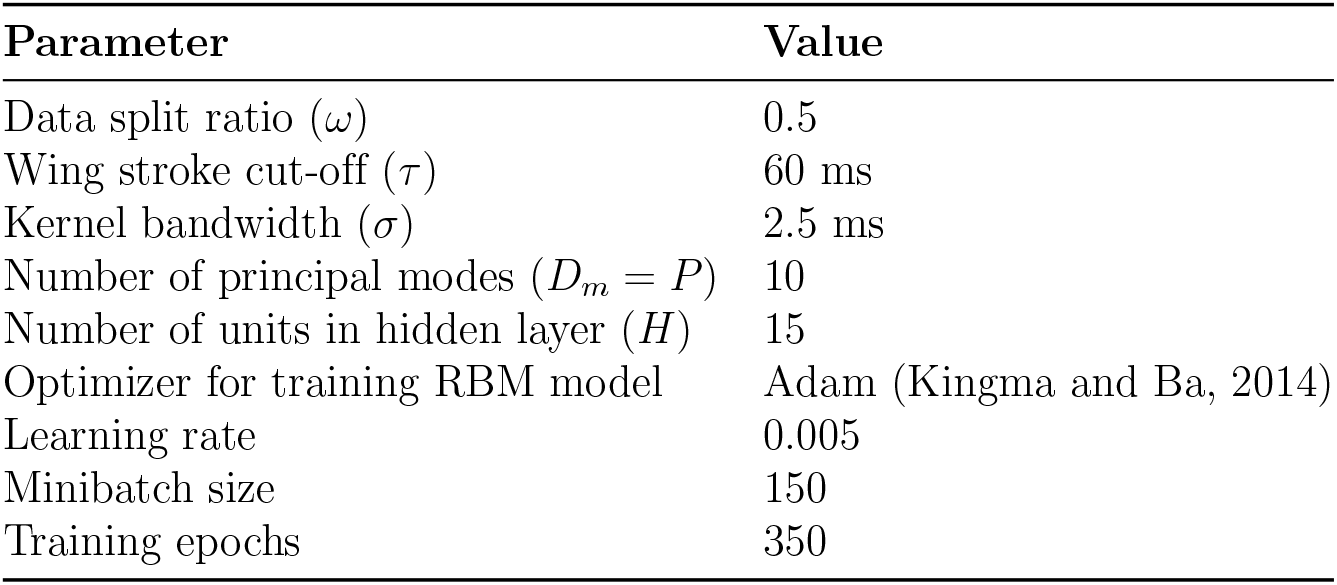
Experimental Parameters.

### 3.3 Results

We used the following values for the free parameters:

In both scenarios, we use the Linear Discriminant Analysis (LDA) for classification which is also applied in Putney et al. (2021) and has shown to perform exceptionally high decoding accuracy. Only one RBM is trained for both scenarios

The results are shown in Figs. 6 and 7. We observe that in both scenarios the performance of the cross-subject neural decoder is bounded between the performances of the first and second benchmarks, with the performance of the subject-specific neural decoder being the upper bound and the performance of the cross-subject neural decoding with no transfer being the lower bound. Note that in Fig. 6 the lower bound varies around 0.16, which corresponds to random choice decoding in our case (as there are 6 stimuli in the experiment, see Section 3.1). The performance of the cross-subject neural decoder is similar in Scenario II but we omit to show it in Fig. 7 to avoid clutter.

**Fig 6:**
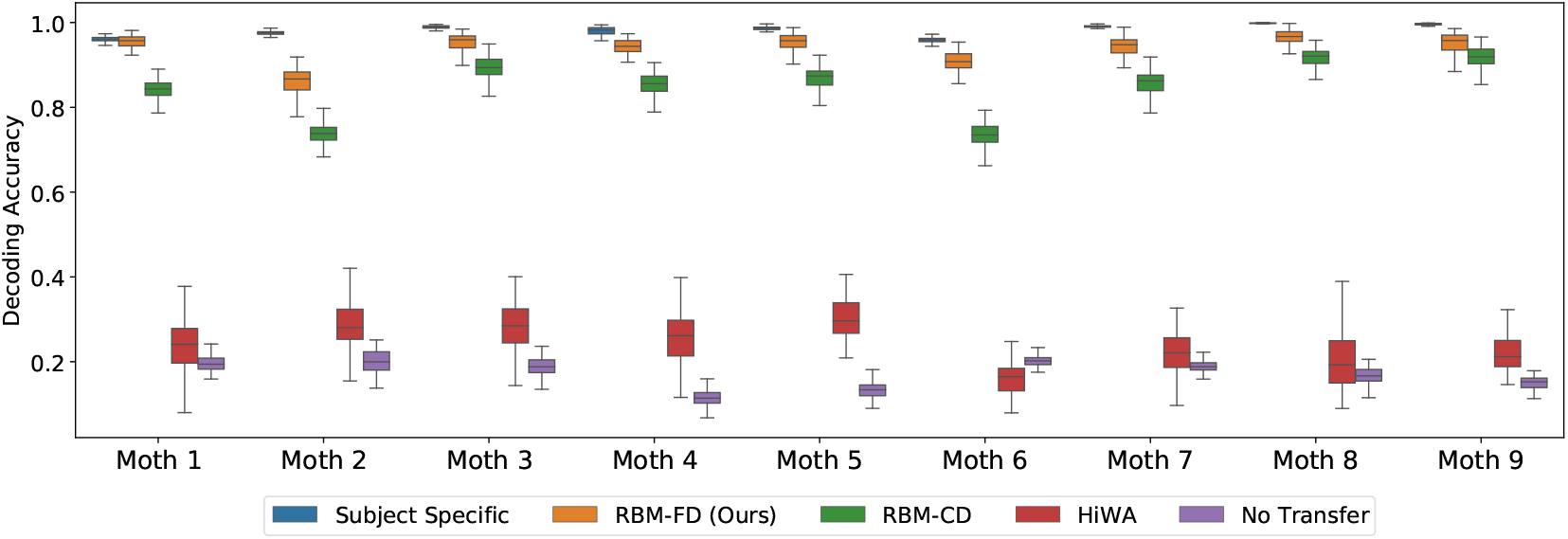
Performance of the cross-subject neural decoder in Scenario I, the error bars are the 95% confidence interval computed using the 100 random initialization (see Fig. 4a and Section 3.2 for details).

**Fig 7:**
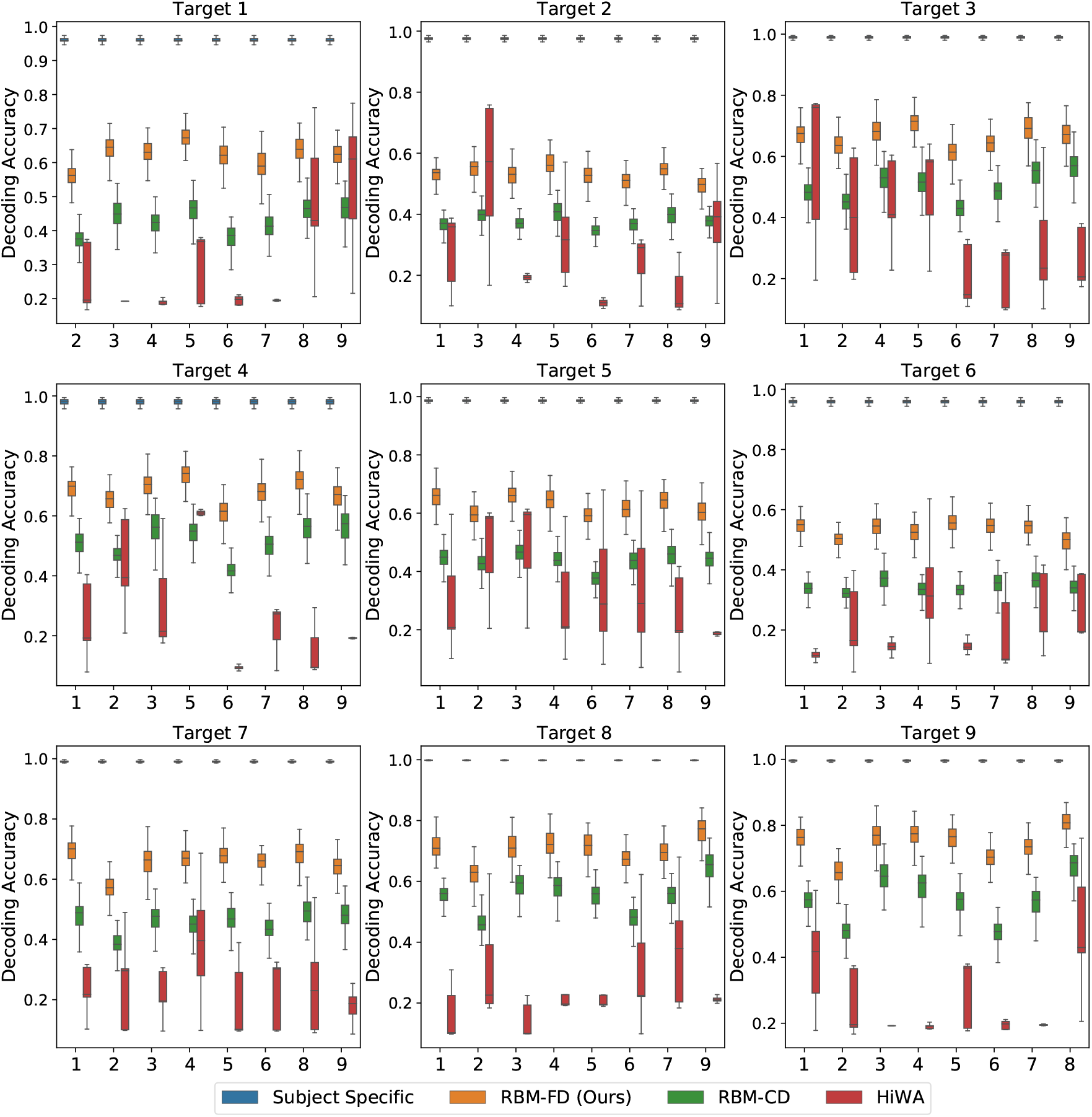
Performance of the cross-subject neural decoder in Scenario II, the error bars are the 95% confidence interval computed using the 100 random initialization (see Fig. 4b and Section 3.2).

We conclude that the performance of the cross-subject neural decoder without transferring the target features to the source is very poor, and produces a poor lower bound, which is also intuitively expected. This can be most easily seen by inspecting Fig. 3 which depicts the feature spaces across the first two modes for each moth. Even though each moth individually exhibits a high degree of class separability (which ultimately results in very reliable subject-specific decoding performance as demonstrated in Fig. 6), there is very little alignment between the geometric distribution of the classes/tasks (i.e., visual conditions) in the space spanned by the first two modes across different moths. In fact, the task-specific representations seem to occupy arbitrary segments of the feature space across the first two dimensions and no discernible pattern can be directly observed; adding even more modes/features which are required to achieve high subject-specific separability, only exacerbates the differences of the feature spaces across moths. As a result, directly decoding any target moth using a neural decoder trained over a different, source moth results in poor performance as reported in Fig. 6.

While HiWA does improve upon the lower bound (cross-subject decoding with no transfer), RBM methods consistently outperform HiWA with higher downstream classification accuracy and smaller variances under different random splitting and initialization in both scenarios. With the complex nature of neural data across subjects, there is no guarantee that different subjects can be mapped through a linear alignment, which could be the reason for HiWA’s limited performance in this dataset. RBM learns a non-linear mapping function that adapts the target to the source feature space where the decoder was trained with few assumptions; in a highly complex setting with multiple subjects and high dimensionality, which is a common theme in neural datasets, our RBM methods could provide a more adaptable alternative to the widely applied alignment methods

The goal of cross-subject transfer learning is to learn a mapping function that takes target features and adapts them to the source feature space where the decoder was trained. We present the RBM-adapted feature space in Fig. 8 which shows the distribution of the target test points in Scenario I after applying the FD-RBM cross-subject transfer with trained RBM model to the corresponding source feature space. Note that the test points are the features coming from a diverse set of targets, namely all remaining (eight) moths; hence, the illustrative result shown in Fig. 8, in addition to the decoding results presented in Fig.6, clearly demonstrate that the FD-RBM model has successfully learned a non-linear transformation that takes a task-specific feature representation from an arbitrary target and maps it into the adequate task-specific region of the source feature space.

**Fig 8:**
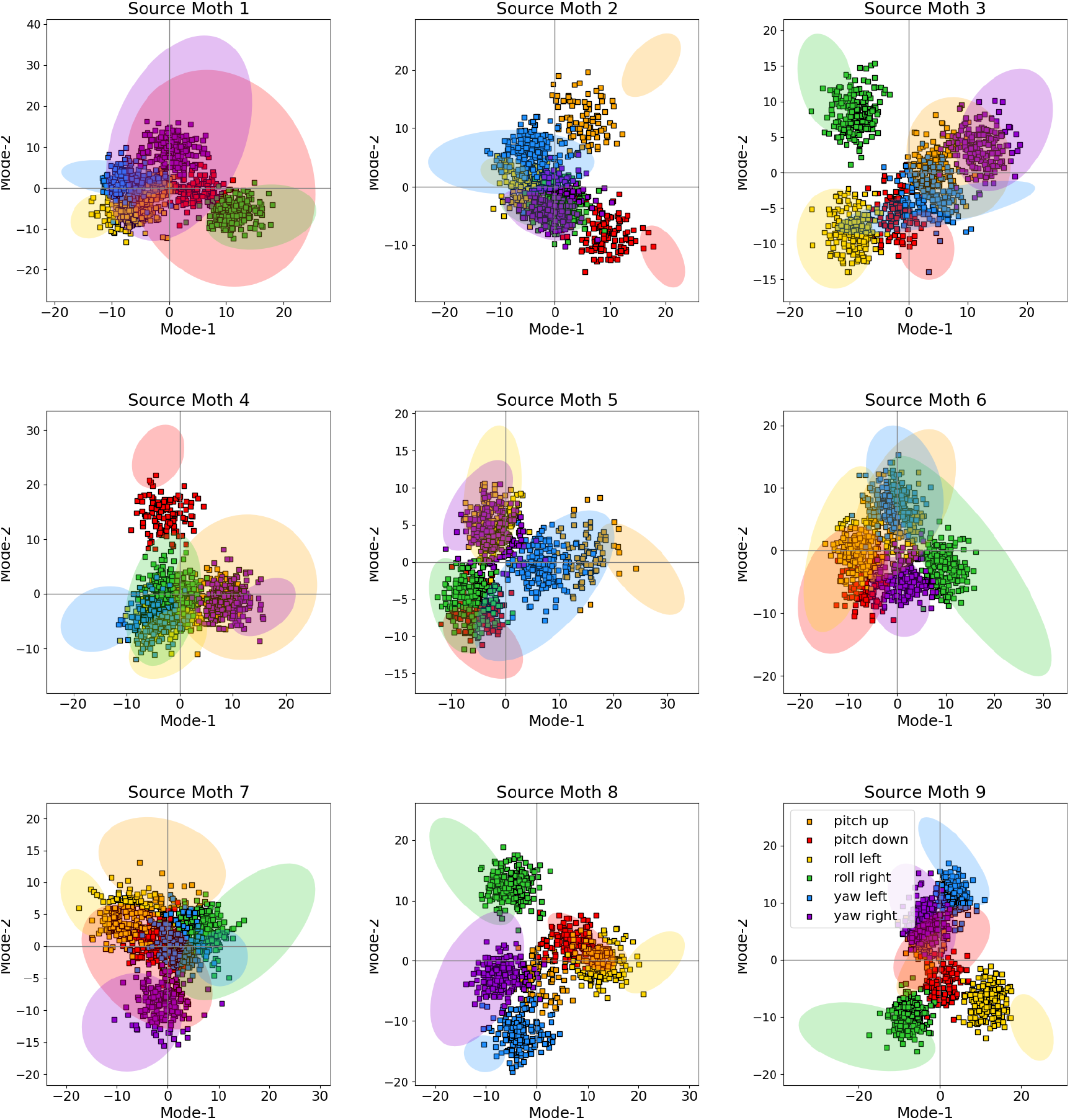
Distribution of target test points (full squares) in the feature spaces of the source moths (represented by the transparent confidence ellipses from Fig. 3) after mapping them using trained FD-RBM model. The mapping corresponds to Scenario I and, as in Fig. 3, only the first two modes of the feature space are shown.

By comparing the results in Fig. 6 with the results in Fig. 7, we also observe that the performance of the neural decoder in Scenario I outperforms the neural decoders in Scenario II. This is an intuitively expected result, as in Scenario II, the size of the joint source feature vector **x**_S_ is *M* − 1 times larger than the target feature vector **x**_D_; that is, in Scenario II we are jointly obtaining the representation of a single target in 8 different source feature spaces. The opposite reasoning applies to Scenario I. Hence, the drop in the performance from Fig. 6 to Fig. 7 is expected. Furthermore, for a given population of subjects indexed in *ℳ*, we can view these two scenarios as the two extreme cases that put the upper (Scenario I) and lower (Scenario II) bounds on the performance.

We also observe that the performance of the cross-subject neural decoder with RBM-FD transfer outperforms the performance of the same cross-subject decoder with RBM-CD transfer. This is an interesting finding, further indicating that in the case of RBMs, the training based on Fisher divergence minimization yields better results in comparison with the more conventional approach based on Maximum Likelihood. This result is also consistent with our findings on popular public datasets such as the MNIST where we used RBM for applications such as compression and reconstruction and where we observed that an RBM trained via Fisher divergence minimization yields higher-quality image reconstructions. In addition to the reliability improvement, we note that training an RBM-FD is less computationally demanding as opposed to RBM-CD which requires Gibbs sampling even during training to obtain estimates for the gradients. However, it should be noted that while the RBM-FD model in general tends to outperform the RBM-CD model, the behavior is ultimately determined by the values of the free parameters, and in the case of the parameters we have selected (listed in the beginning of this section), the above observations are valid.

## 4 Discussion

The design of reliable, robust, and computationally efficient solutions for cross-subject mapping and combining data across subjects is a challenging problem in neuroscience. In this paper, we proposed a novel framework for learning the joint distribution of target and source feature representations across a set of subjects using the undirected graphical model and the flexible non-linear generative model RBM. To improve the efficiency of traditional RBM, we considered an alternative training method for the RBM that minimizes the Fisher divergence and allows the gradient to be computed in closed form, alleviating the need for Gibbs sampling during training. We evaluated our proposed scheme against recently developed distribution alignment methods, HiWA, on a neural decoding task and experimental data collected from nine hawk moths during a comprehensive motor program where the moths are subject to a total of six visual stimuli, and we found substantial improvement over HiWA in two transfer scenarios. The results verified the viability of the solution. These approaches, which are highly flexible with few assumptions, show promise in generalizing features of complex neural datasets across individuals, tuning neural interfaces to subject-specific features, and leveraging data across multiple subjects when experiments are limited in time or completeness.

### Application of Restricted Boltzmann Machine in Neuroscience

Restricted Boltzmann machines are widely applied as a tool to study neuroscience problems. The bipartite network structure has long attracted neural science interests to study the structural con-nection and functional activites in the brain (Hjelm et al., 2014). Recently, RBM has been applied to study the structural connectivity of zebrafish larvae and identified coarse-grained neural grouping (van der Plas et al., 2023). Combining a recurrent element with the RBM for temporal analysis, a variant of RBM, the recurrent temporal RBM was shown to capture the temporal dynamic of the whole brain activities of zebrafish larvae (Monnens et al., 2024).

Our work considers a different application in neural science, where we leverage the undirected graph structure of the RBM to learn joint distributions of neural activities, and then utilize this joint distribution to map between multiple subjects for independent downstream tasks. In general, with its special bipartite network construction and its undirected probabilistic graphical model structure, RBM is a highly flexible tool to further explore neural science for understanding brain structures, and dynamical neural activities, or apply to cross subject mapping.

### Efficient and versatile cross-subject mapping

Many cross-subject mapping methods have been proposed to improve cross-subject mapping methods to be more accurate, robust, and versatile. One way to map between subjects is through alignment, which aligns the neural representation between two subjects directly after simple preprocessing such as PCA (Gallego et al., 2020; Lee et al., 2019); one can also map the neural representation to a latent manifold before learning a mapping between the source and the target (Dabagia et al., 2023; Degenhart et al., 2020; Herrero-Vidal et al., 2021). More recently, generative algorithms emerging from deep learning were also applied to cross-subject mapping. For example, a Generative Adversarial Network was applied to transfer human activities between two subjects (Soleimani and Nazerfard, 2021) while a conditional VAE was applied to transfer latent neural representation between monkeys (Angjelichinoski, Pesaran and Tarokh, 2020).

However limited studies focus on improving the general efficiency of cross-subject mapping methods, especially when multiple subjects are present in a complete dataset. Most of the cross-subject mapping methods deal with two subjects where at most two transformations are required to enable cross-subject mapping (Herrero-Vidal et al., 2021; Lee et al., 2019). As the number of subjects grows, the number of models needed to transfer between all subjects grows exponentially. To reduce the number of mappings when multiple subjects exist, researchers have developed a feature transfer that needs to be learned in a supervised manner (Zhao et al., 2021). Our work approaches a similar multi-subject/multi-target mapping problem from a probabilistic standpoint: by using RBM and estimating the joint distribution of all subjects at the same time, we learn the conditional probability that allows the mapping of an arbitrary number of targets to an arbitrary number of sources. This process improves the efficiency of cross-subject mapping with no required labels as we directly learn the distribution of features without a decoder.

Cross-subject mapping in a generative, probabilistic way through RBM also allows the RBM to be more versatile to apply in various settings. A common drawback of current methods of cross-subject mapping requires the availability of the source subject’s data during test time. For example, the alignment-based algorithms can only correctly compute the alignment matrix when both the source and test are available (Degenhart et al., 2020; Lee et al., 2019). However, the RBM allows us to transfer different target subjects to the space of the source subject without additional knowledge regarding the source beyond its training data. This is enabled by the probabilistic modeling of RBM. With the joint distribution of the subjects represented by the RBM, conditional distribution can be easily computed for unknown subjects, allowing cross-subject mapping when a subject is not available. The probabilistic framework is also applied to align multiple subjects’ neural responses for training a pooled decoder to classify odor stimuli (Herrero-Vidal et al., 2021).

### Limitations

While the RBM architecture with fisher divergence proposed in this study greatly improves the efficiency of cross-subject mapping over many subjects, limitations remain from several perspectives: its asymmetrical transformation, limited extensions to quantify similarities between subjects, and its lack of structure in the hidden layer that removes interpretability from this architecture.

Many cross-subject mapping algorithms are symmetrical, meaning that transforming the one subject to another subject can be directly applied to transform in the opposite directions with limited changes to the transformation function. Usually, the symmetric methods map two subjects onto a shared embedding space, which is widely applied in neural alignment methods such as (Harvey et al., 2024; Kriegeskorte et al., 2008). The RBM cross-subject mapping method is directional: the transformation from one subject to all of the other subjects learned by an RBM is characterized through complex neural networks that are non-linear. Directional transformations inevitably will make the transformation process less efficient, but the capability of mapping between multiple subjects at once compensates for the lack of symmetry in the transformation.

Another drawback of our RBM methods is that we cannot easily derive a distance or similarity metric to quantify the similarities between two (groups) of subjects. Many alignment-based methods, such as Harvey et al. (2024), can directly extract the similarity information between two subjects once the alignment is complete. We refer to Klabunde et al. (2023) for a more comprehensive review of similarity metrics derived from cross-subject mapping or alignment. With RBM being an estimator of the joint distribution of all subjects capable of sampling from conditional distributions, it is possible to compute information-based metrics such as the conditional entropy between two (groups of) subjects. One issue with conditional entropy, however, is that this metric is not symmetric, and thus cannot be used explicitly to quantify similarities between subjects. Symmetrical information metrics like the mutual information between two (groups) of subjects cannot be computed with the current architecture as marginal distribution of subjects is required for that computation. However, the conditional entropy is not strictly a distance metric because it is not symmetric. Modifying the current RBM architecture could lead to a system that can quantify similarities between subjects through mutual information, which is symmetric, but that is beyond the scope of this current study.

The lack of interpretability is another limitation of our current architecture. Cross-subject mapping methods that utilize algebraic solutions can identify how each feature of the target subject is mapped to the space of the source subject. For example, the Procrustes Alignment method computes a simple transformation matrix that maps the target data onto source space (Harvey et al., 2024). However, it is difficult to understand the operations applied to the target data by the RBM to transform them into the source space due to their non-linear nature. Various network analysis methods could be used to test the sensitivity of the representation and decoding to specific inputs (Yang et al., 2022). So where the goal of an analysis is prediction, classification, or other types of questions amenable to non-linear ML approaches, our RBM with Fisher divergence method provides a flexible, efficient framework for combining neural data across subjects even when the recordings are sparse spiking representations

### Future work

Several extensions of the presented approach are possible and are currently part of our ongoing work. First, the derivation of the conditional distribution of the source features given the target features from the joint distribution represented by an RBM is a direction worthwhile pursuing as it relates to the cross-subject mapping problem and would avoid the issues related to noise-like initialization in the approach presented in this paper. Second, learning subject-invariant RBM models that can also predict feature representations of unseen target subjects is a key direction that should be pursued since it is directly related to the generalization capability of the approach. Finally, we are also investigating the application of the model to other neural signal modalities, including non-invasive modalities such as EEG signals.

## Footnotes

1 We omit the detailed derivation of the gradients for brevity, and we note that the gradients can be derived through straightforward application of matrix calculus.

